# The Evolution of Primate Body Size: Left-skewness, Maximum Size, and Cope’s Rule

**DOI:** 10.1101/092866

**Authors:** Richard C. Tillquist, Lauren G. Shoemaker, Kevin Bracy Knight, Aaron Clauset

## Abstract

Body size is a key physiological, ecological, and evolutionary characteristic of species. Within most major clades, body size distributions follow a right-skewed pattern where most species are relatively small while a few are orders of magnitude larger than the median size. Using a novel database of 742 extant and extinct primate species’ sizes over the past 66 million years, we find that primates exhibit the opposite pattern: a left-skewed distribution. We investigate the long-term evolution of this distribution, first showing that the initial size radiation is consistent with plesiadapiformes (an extinct group with an uncertain ancestral relationship to primates) being ancestral to modern primates. We calculate the strength of Cope’s Rule, showing an initial tendency for descendants to increase in size relative to ancestors until the trend reverses 40 million years ago. We explore when the primate size distribution becomes left-skewed and study correlations between body size patterns and climactic trends, showing that across Old and New World radiations the body size distribution initially exhibits a right-skewed pattern. Left-skewness emerged early in Old World primates in a manner consistent with a previously unidentified possible maximum body size, which may be mechanistically related to primates’ encephalization and complex social groups.

## I. INTRODUCTION

Primates are a taxonomically and geographically diverse clade with hundreds of extant and extinct species found across South and Central America, Africa, and Asia, and with extinct species found in North America and Europe. Originating over 66 million years ago, primates exhibit a number of unusual behaviors, including complex, hierarchical social orders (Van Schaik 1996; Van Schaik and Van Hooff 1983), intergroup aggression and warfare (Manson et al. 1991; Mitani et al. 2010), tool-use in a variety of manners (Breuer et al. 2005; Ottoni and Izar 2008), and both arboreal and terrestrial lifestyles. They are perhaps best known for their relatively large brain size compared to their body size (Boddy et al. 2012; Shultz and Dunbar 2010), as measured by their encephalization quotient. This large brain size is believed to play a fundamental role in their complex social behavior. Extant primates tend to exhibit a high encephalization quotient, despite a broad distribution of body sizes across the clade, ranging from the pygmy mouse lemur (55 g) to the gorilla (130, 000 g).

Here, we investigate the evolution of primate body sizes over the past 66 million years. We study how the body mass distribution of primate species varies across both time and changing ecological conditions, and how these patterns contrast with those of terrestrial mammals in general. We focus on body size as it is a fundamental variable for a species, is relatively easy to measure, and is comparable across extant and extinct species. Moreover, body size is closely related to many key ecological traits, including habitat, diet, geographical location, life span, population size (Brown 1995; Cardillo et al. 2005; Smith et al. 2002; West et al. 2002*a*), population growth (Savage et al. 2004), evolutionary fitness (Brown et al. 1993), and extinction risk (Tomiya 2013), as well as physiological and evolutionary characteristics, such as metabolic rate (Gillooly et al. 2001; Kleiber 1947), DNA nucleotide substitution rate (Martin and Palumbi 1993), brain size (Kappelman 1996; Lande 1979; Martin and Harvey 1985), and skeletal measurements (Gingerich et al. 1982; Jungers 1985). This makes body size a convenient proxy variable (Lucas et al. 2008) that relates generally to important ecological, physiological, and evolutionary characteristics of species, and an ideal characteristic for a comparative focus within a given clade (Gittleman 1985). A deeper understanding of primate body size patterns will shed new light on the evolutionary history of this important clade, particularly relative to the concurrent expansion of terrestrial mammals and their sizes. The lens of species body distributions also provide a novel approach for characterizing the unusual evolutionary trajectory of primates, which produced high encephalization, sophisticated cognitive functions, complex social structures, and humans (Dunbar and Shultz 2007; Shultz and Dunbar 2007).

The extant and extinct body mass distributions of terrestrial mammals as a group, as well as those of subclades such as horses (Shoemaker and Clauset 2014) and whales (Clauset 2013), are known to exhibit a right-skewed pattern over multiple orders of magnitude, indicating relatively fewer large species than small species (Clauset and Erwin 2008; Kozłowski and Gawelczyk 2002; McShea 1994; Smith and Lyons 2011). This canonical pattern also appears in other major animal clades, including birds, fish, and insects (Kozłowski and Gawelczyk 2002). The shape of these distributions is a robust outcome from the interplay of macroecological and evolutionary processes under physiological constraints (Clauset and Erwin 2008). Within an evolving clade, the body masses of its species follow a constrained, cladogenetic (branching) random-walk through time. The first constraint on the evolution of body size is a minimum viable body mass, which is dictated by thermoregulation, metabolic rate, and the clade’s climactic conditions (West et al. 2002*b*). The maximum achievable species size is determined by a crossover point, when the risk of extinction, which grows with body size as a result of smaller population sizes and demographic stochasticity, exceeds the speciation rate (Clauset 2013). Between these minimum and maximum sizes, the right-skewed pattern is the outcome of an evolutionary trade off between the short-term advantages of increased size and the long-term risks of increased extinction rates for lineages within the clade. This pattern does not require a bias toward larger sizes within a lineage, as in Cope’s rule (Alroy 1998), but may be enhanced by one.

However, it is unclear whether the right-skewed pattern occurs across all mammalian sub-clades, and whether the same underlying processes govern the evolution of body masses for primates, given their unique evolutionary trajectory, social structure, and unusual characteristics. Additionally, primates provide an unusual opportunity to study the evolution of body mass within a single clade, but across environmental conditions, as primates have two separate radiation events separated in time by 36 million years and in space between the Old and New Worlds. Old World primates originated approximately 66 mya in Asia (Beard 2004) while New World primates originated roughly 30 mya when small groups of primates arrived in the Americas. The temporal and spatial separation of these two radiation events of 36 million years and two distinct landmasses provides a natural experiment through which we may compare the radiation of a single clade under distinct climatic and ecological constraints, further illuminating the roles these variables can play in the evolution of body size.

Using a novel database of species-level primate body masses and first and last appearance estimates covering the past 66 million years, we study primate radiations, the overall evolution of primate body mass through time, and their relationship to environmental characteristics. We compare the evolution of primate body mass with that of terrestrial mammals to gain insight into the relationship between primate evolutionary history and the enclosing clade of all terrestrial mammals. We additionally investigate the relationship between primates and plesiadapiformes (an extinct order either closely related to primates or potentially the predecessor of primates (Silcox 2007)) by examining the body size dynamics within the initial radiation of primates with and without including plesiadapiformes as direct ancestors. We then explore how the evolution of primate body mass correlates with temperature and other environmental factors, such as the radiation of grasslands. Finally, we consider the generality of these insights through a direct comparison of size-related evolutionary patterns in the Old and New World primate radiations.

## II. METHODS

### A. Data Collection and Mass Estimation

To study the distribution of primate body size and its evolution over time, we constructed a novel database consisting of 1024 primate and 86 plesiadapiform species belonging to 373 genera (Alroy et al. 2015; Beard 1987, 1988; Beard and Houde 1989; Bloch et al. 2002, 2001, 1998; Boubli and Ditchfield 2001; Bown 1982; Bown and Rose 1976, 1984, 1987, 1990, 1991; Burger 2007; Cameron 2004; Chopra 1978; Ciochon and Fleagle 1987; Covert and Williams 1991; Cuozzo 2008; Davidson 1987; Di Fiore et al. 2015; Eaton 1982, 1985; Emry 1990; Fleagle 2013; Fox 1984, 1990, 1991, 2002; Fox and Scott 2011; Fox et al. 2010; Frost et al. 2015, 2003; Gazin 1942, 1958, 1962, 1968, 1969, 1971; Gingerich 1975, 1976, 1989, 1993, 1995; Gingerich and Dorr Jr 1979; Gingerich and Haskin 1981; Gingerich et al. 1983; Gingerich and Simons 1977; Godinot and Mahboubi 1992; Gunnell 1985, 1989, 1992, 1995*a*,*b*, 1998, 2002; Gunnell and Gingerich 1981; Hartman 1986; Holroyd and Strait 2008; Honey 1990; Hunter et al. 1997; Hunter and Pearson 1996; Jones et al. 2009; Kelly and Whistler 1994; Kihm 1992, 1984; King et al. 1999; Kirk and Williams 2011; Krause 1978; Krause and Gingerich 1983; Krishtalka 1978; Krishtalka et al. 1975; Lillegraven 1980; MacPhee and Iturralde-Vinent 1995; Mason 1990; McGrew and Patterson 1962; McGrew and Sullivan 1971; McKenna 1960, 1990; Mootnick and Groves 2005; Muldoon and Gunnell 2002; Murphey and Dunn 2009; Qi et al. 2006; Rasmussen et al. 1995; Rasmussen 1996; Rasmussen and Nekaris 1998; Rigby 1980; Robinson 1968, 1994; Rose 1981, 1995; Rose et al. 1993; Rose and Bown 1982, 1991, 1996; Rose and Gingerich 1976; Rose et al. 1999; Schiebout 1974; Scott 2003; Scott and Redman 2016; Scott and Fox 2005; Shigehara et al. 2002; Silcox 2001; Simons 1961; Simpson 1935, 1936, 1937; Stirton 1951; Stirton and Savage 1949; Stock 1934; Storer 1990; Swindler 2002; Szalay 1969, 1976; Takai and Anaya 1996; Tejedor et al. 2008; Van Valen 1994; West 1973, 2015; Williams and Kirk 2008; Williams et al. 2007; Winterfeld 1982; Wood et al. 1979; Zinner et al. 2013; Zonneveld et al. 2000*a*,*b*). We included all primate and plesiadapiform species with published first and last appearance estimates or estimated body masses, yielding a total of 742 species with all three estimates (Table S1). These 742 species were used for all subsequent analyses.

A species list of extinct primates and plesiadapiformes was first constructed from the data recorded in Fossilworks (Alroy et al. 2015), an online and community-supported repository of fossil data. For each species, we recorded first and last appearances from primary literature or Fossilworks (Alroy et al. 2015). If no detailed first and last appearance estimates were available for a given species, we used the estimated epoch time periods (Fleagle 2013). Genus-level first and last appearance dates were not utilized unless only a single species in the genus is currently known.

We estimated species body mass using multiple methods. When available, we included estimates of body masses from the primary literature. Following common body mass estimation methods (Gingerich et al. 1982), we additionally used allometric scaling to estimate body mass from dental measurements for the first, second, and third upper and lower molars in addition to the fourth upper and lower premolar, providing a total of 16 possible measurements for each species. We recorded mesiodistal lengths, buccolingual widths, and for teeth without buccolingual measurements, we preferred anterior breadth measurements to posterior breadth. We also preferred trigon breadth to talon breadth measurements when available. We used allometric scaling to then estimate body mass from each dental measurement from a total of 16 models of the form log(*M*) = *α* log(*D*) + *β* where *M* is the estimated body mass in grams, *D* is the dental measurement for the particular model, and *α* and *β* are the slope and intercept of the best fit (least squares) line through the transformed data. All models produced *R*^2^ values between 0.73 and 0.91 (model details are included in Appendix S1). When multiple estimates were available, we averaged estimates from all applicable molar models and previous literature estimates to obtain a single species mass estimate.

All extant species with mass estimates from *Handbook of the Mammals of the World — Volume 3: Primates* (Zinner et al. 2013) or PanTHERIA (Jones et al. 2009) were included in our database, yielding mass data on 498 extant primate species. For extant species with unknown first appearance dates, we show first appearance dates as 1 Ma for visualization purposes only. For both extinct and extant species, when multiple subspecies were listed, we averaged the mass of each sub-species to obtain a single species-level estimate. Similarly, we took the average mass when a minimum and maximum were included for a single species or when male and females masses were presented separately, yielding a single point estimate of body mass per species.

### B. Data Analysis

Dividing the past 70 million years into 5 million year bins, we calculated the mean, minimum, maximum, and skewness of the primate body size distribution in each bin as summary statistics. A bin size of 5 million years provides a balance between ensuring a sufficient number of points within each window, avoiding statistically anomalous changes in the summary statistics caused by small sample size, and sudden changes that are consequences of approximate first and last appearance dates and not evolutionary mechanisms. This approach is consistent with previous methods (Clauset and Erwin 2008; Clauset and Redner 2009; Shoemaker and Clauset 2014), and will facilitate a direct comparison across studies. To calculate these summary statistics, we include all species that existed in each time bin, regardless of their first and last appearances. Thus, long-lived species or species that are present in multiple bins are included in all time bins that they overlap. The mean, minimum, and maximum body size provide summary estimates of how the body size distribution shifts through time, and facilitate a test of whether the extremes of the distribution follow a pattern similar to the central tendencies. Skewness is a measure of a distribution’s asymmetry, and a positively skewed, or right-tailed, distribution exhibits relatively fewer very large species versus very small ones (Clauset and Erwin 2008). To provide a comparison with terrestrial mammals as a whole, we calculated the skewness for the body sizes of North American mammals using data from 2008. Finally, we also calculated all four summary statistics for all extant primate species.

To estimate the average change in body mass between descendants and ancestors, we use Alroy’s within-lineage definition of Cope’s rule (Alroy 1998) to calculate the strength of Cope’s rule for species in every 5 million year window and for extant species. To do so, we first defined each species in a given time bin as a descendant. We then selected, uniformly at random from the set of plausible ancestors, one species to assign as its ancestor. Plausible ancestors are defined as all species in the descendant’s genus for whom the descendant’s first appearance date falls on or between the ancestor’s first and last appearance dates. If no such species within the genus is currently known, we instead assigned as the ancestor the species within the genus with the last appearance date closest to the first appearance date of the descendant. When a descendant species had no plausible ancestors and there was no within genus species with a last appearance date coming before the first appearance of the descendant, a plausible ancestor-descendant pair cannot be determined. In this case, the species was ignored when calculating the strength of Cope’s rule. Once all ancestor-descendant pairs are determined, we calculate the pairwise change in sizes as ln(*M_d_/M_a_*) where *M_d_* is the mass of the descendant species and *M_a_* is the mass of the ancestor. The average of this quantity over all descendant species and over 100 trials is a maximum likelihood estimator of the average strength of Cope’s rule for that time bin (Alroy 1998). Following the same procedure for extant species provides an estimate of the strength of Cope’s rule in the most recent time period.

We estimated “radiation cones,” which represent the trajectory of the minimum and maximum species body masses during the initial radiation of a clade. This allowed us to compare both the size-range expansion rates of Old World versus New World primate radiations and the expansion rates of primates versus plesiadapiforms, which radiated 3 million years prior to the first known appearance of primates. These radiation cones begin with all known species at the beginning of the radiation (furthest date from present) and expand along the trajectories of the minimum and maximum body masses of the radiating clade. Calculated radiation cones measure time, *t,* as millions of years since 70 million years ago. The radiation cone for plesiadapiformes is defined by the lines log_10_(*M*) = −0.09*t* + 2.15 (minimum mass) and log_10_(*M*) = 0.15*t* + 1.84 (maximum mass) spanning 66 to 55 mya. The radiation cone for Old World primates is defined by the lines log_10_(*M*) = −0.07*t* + 1.90 (minimum mass) and log_10_(*M*) = 0.13*t* + 2.09 (maximum mass) spanning 63.25 to 55 mya. Finally, the New World expansion of primates spans from 23 mya to present day and is bounded between log_10_(*M*) = −0.01*t* + 3.30 (minimum mass) and log_10_(*M*) = 0.04*t* + 1.51 (maximum mass).

Extinct and extant New World and Old World primate body size distributions are smoothed using a normally distributed kernel density smoother (Wasserman 2002). Climate conditions from 66 mya to present are used to compare body size changes through time to key climate changes. All climate data is from Zachos et al. (2001).

## III. RESULTS AND DISCUSSION

Across terrestrial mammals, birds, lizards, and mammalian subclades such as Equidae (horses) and cetaceans (whales, dolphins, and porpoises), extant body mass exhibits a canonical right-skewed distribution (Kozłowski and Gawelczyk 2002). Surprisingly, in contrast to this common pattern, we find that extant primates exhibit a left-skewed distribution of body sizes (skew = −0.27 ± 0.11) (Figure 1). That is, primates exhibit the opposite pattern relative to what is typical for mammals (Clauset and Erwin 2008), with relatively fewer small primates than large primates.

**FIG. 1.**
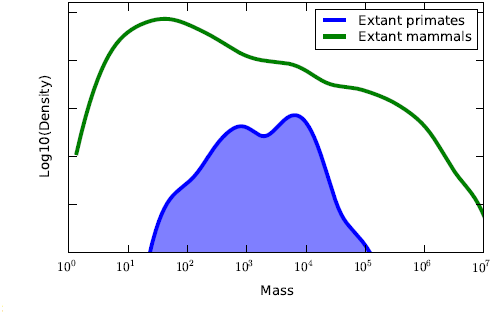
Extant primate (blue online; dark grey print) and terrestrial mammal (green online; light grey print) body size distributions.

We further explore the evolution of the primate body size distribution and its origins to better understand the evolutionary pressures responsible for the left-skewed pattern exhibited by extant primates. As with other mammals, fossils from the Paleocene describing primate ancestors largely consist of teeth, small intact bones, and bone fragments (Fox and Scott 2011; Hunter and Pearson 1996; Silcox 2007), leaving out critical features used in trait-based phylogenies, e.g., post orbital bar, flattened nail on hallux (Wible and Covert 1987). However, as the fossil record and molecular phylogenies both improve, evidence increasingly supports defining Euprimates (also called true primates) as originating either in the very late Cretaceous or early Paleocene (65 — 62 Ma), closely matching our estimate of primate first appearance of 63.3 mya. Several species of early primates in the genus *Purgatorius* have been identified from Paleocene beds in Montana, USA and Saskatchewan, Canada (Fox and Scott 2011; Van Valen and Sloan 1965). However, diversity of *Purgatorius* species in these beds suggests that their radiation likely began in the late Cretaceous (Scott and Redman 2016). These species are now generally accepted as the likely earliest common ancestors of modern primates and are placed in the order plesiadapiformes (Bloch et al. 2007). Yet, plesiadapiformes are often treated as a paraphyletic clade excluding modern primates, although fossils discovered in recent decades show plesiadapiformes had many of the traits considered exclusive to Euprimates, e.g., grasping hallux with nail and petrosal bulla in skull (Bloch et al. 2007), indicating that they may be early common ancestors of primates.

Our analysis of early body size radiations further support the hypothesis that plesiadapiformes are early ancestors of primates. Our earliest first appearance dates collected for primates show four species *(Palenochtha weis-sae, Picrodus calgariensis, Edworthia lerbekmoi,* and *Torrejonia wilsoni*) originating 63.3 mya and characterized by body sizes spanning two orders of magnitude (Figure 2). This substantial size disparity suggests that the primate clade actually originated earlier in time. Obviously, it would be impossible for more than one species, let alone several species spanning multiple orders in body size, to begin a new clade. One of these species could be the common ancestor of the others, though this is unlikely as it would require an extremely rapid divergence in species sizes within the newly radiating clade in order to match the empirically observed size range. However, the radiation cone of plesiadapiformes from 66 to 55 mya is similar to that of primates both in terms of the rates of size expansion (0.24 vs 0.20; Figure 2, difference in slopes of dashed lines) and the estimated initial founder body size (114 g vs 112 g) (Figure 2, dashed lines, extrapolated), providing support to the hypothesis that plesiadapiformes are likely early common ancestors of modern primates. In particular, the difference in the expansion rates of plediadapiformes and primates may be interpreted as a percentage increase in body size per million years.

**FIG. 2.**
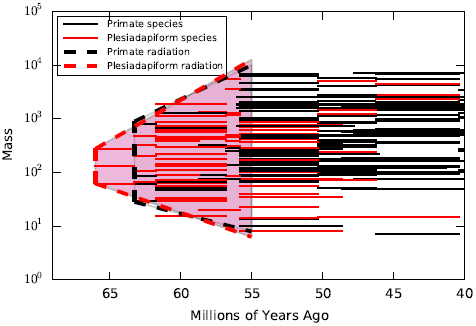
Body sizes of primate (black) and plesiadapiform (red online; grey print) species at the initial radiation event. Solid lines are individual species and dashed lines depict radiation cones (see text) with (red online; grey print) and without plesiadapiformes (black).

The strength of Cope’s rule may also be interpreted as an average percent change in body size over time. Here, the rate difference, 0.04, falls well within a single standard deviation of our estimates for the strength of Cope’s rule in the 60–55 mya and 55–50 mya windows (0.035±0.009 and 0.033±0.008 respectively). This suggests that the observed rate difference is a product of noise as opposed to a true difference in the expansion rates of plediadapiformes and primates. Additionally the three earliest plesiadapiromes *(Purgatorius titusi, Purgatorius unio*, and *Pandemonium dis*) exhibit intermediate body sizes between the minimum and maximum of the earliest primates, and span only a single order of magnitude, further supporting the notion that the group plesiadiformes should be included within the Euprimates.

After origination, the size distribution of primates expands at both the upper and lower extremes from 66 to 55 mya, at which point it exhibits a minimum size of 8 g and a maximum size of 7000 g (Figure 3a,b). This range in body sizes represents the maximum disparity (nearly a factor of 900) observed over the past 66 million years. Both maximum and minimum primate size then remain relatively stable until the observed minimum body size increases to 50 g around 40 mya. However, the upper extreme of the distribution remains fairly stable, at 7000 g, until 23 mya, after which it again climbs over time to a maximum of 225, 000 g *(Gigantopithecus blacki*) at 2.58 mya.

**FIG. 3.**
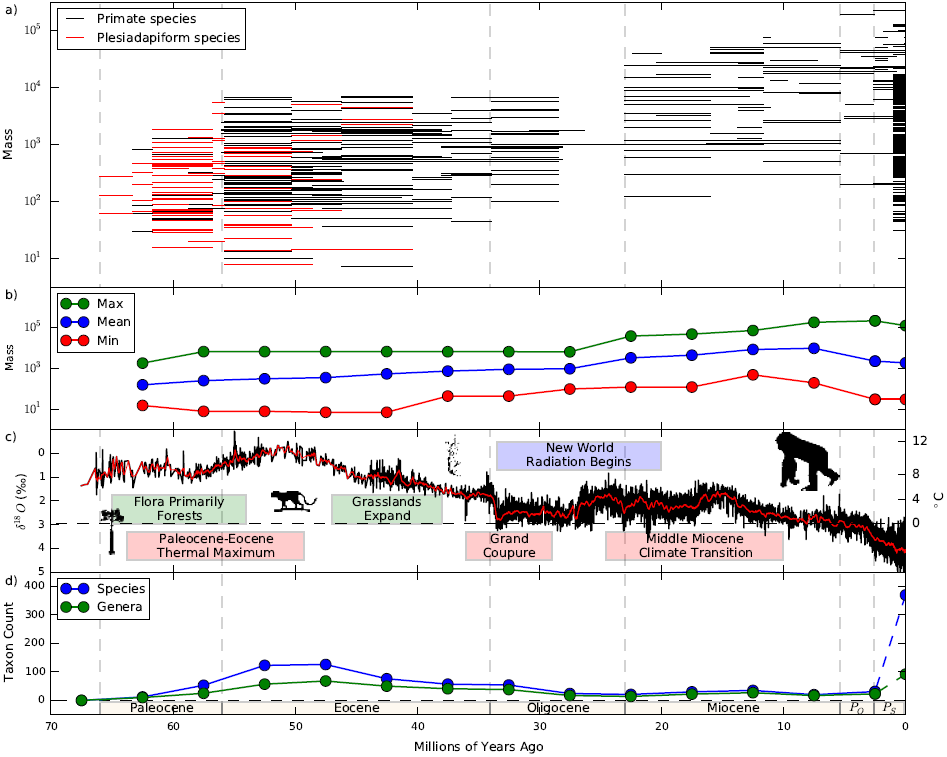
The evolution of primate body sizes over 66 million years (*P_O_* and *P_S_* denote the Pliocene and Pleistocene, respectively). a) Body sizes of primate and plesiadapiform species, with each line spanning the first and last appearance of the given species. b) Minimum, mean, and maximum body sizes, at 5 million year intervals. c) *δ*^18^*O* measurements and corresponding global temperature estimates (data from (Zachos et al. 2001)), along with key ecological features. d) Empirical taxonomic diversity among primates, at both species and genus levels.

Notably, there is a pronounced fall in taxonomic diversity in our database from 23 to 28 mya, which coincides with the well known late Oligocene gap in the primate fossil record (Tavaré et al. 2002). This gap occurs soon after the Grande Coupure extinction event and coincides with the beginning of New World primate radiation. However, our data suggest that this apparent decrease in primate diversity is an artifact of the fossil record. If the Oligocene primate fossil record gap represented a true primate extinction event, we would expect to observe both decreased taxonomic diversity and reduced size disparity immediately following the extinction event, and then possibly a steady expansion (likely at a similar rate as the original radiation) of one or the other or both. Instead, immediately following the decrease, at the beginning of the Miocene, we observe both large taxonomic diversity and large mass disparity, spanning 2.5 orders of magnitude. Following this gap in the fossil record, the minimum size remains relatively stable until present day, while the maximum observed size increases slightly in both the Pliocene and Pleistocene.

Our database represents a relatively high and consistent sampling across time (Figure 3d), implying that trends in body sizes and diversity are likely not artifacts of variability in data coverage or sampling effort.

Environmental conditions and global temperatures are expected to be substantial sources of macoevolutionary pressures that, in turn, influence long-term patterns of body size across species (Angilletta et al. 2004). As such, changes in the environment can have considerable impact on the evolution and distribution of body sizes within a clade. For example, we may expect a decrease in temperature to correspond to an increase in average body size (Bergmann’s rule (Bergmann 1848)), although many exceptions to this general trend exist and the trend is known to be plastic (Angilletta et al. 2004; Arendt 2011). Changes in measured delta oxygen 18 (*δ*^18^*0*) levels, which represent the ratio of stable isotopes oxygen-18 and oxygen-16, and correspond to changes in global temperature (Zachos et al. 2001) over the past 66 million years (Figure 3*c*) indicate a reliable correlation between global temperatures and body size. Specifically, maximum and mean mass increase during the initial primate radiation, which corresponds to the increased temperature leading to the Paleocene-Eocene thermal maximum. Then, the maximum size remains constant while the minimum size increases as global temperatures decrease into the Oligocene. After the middle Miocene climate transition, there is an increase in maximum, mean, and minimum primate body size as global temperatures decrease. Although this correlation is consistent with what we might expect from Bergmann’s rule, we emphasize that it is only a correlation. It remains unclear what mechanistic role, if any, global temperatures and thermoregulation might have played in these changes.

The expansion of grasslands during the Eocene also coincide with a large jump in the minimum size of primates, a period during which smaller arboreal primates become less common and primates begin using more open habitats and spending more time in terrestrial environments. A reduction in access to, and dependence upon, trees alters food abundance and availability, habitat range, and predator avoidance strategies in such a way that larger body sizes are expected to be advantageous. In comparison, an environment with little open space and relatively dense trees should tend to favor smaller, more arboreal species, whose weight and caloric needs are more easily supported by trees.

While transitions in primate minimum and maximum sizes occurred over relatively short time periods, changes in overall body size distributions occurred more gradually (Figure 4). Across primate evolution, body size distributions gradually shift from right-skewed to left-skewed as the distribution evolves toward larger sizes, with increases in both the minimum and maximum sizes of primates. This shift is gradual through time (Figure 4 comparison of black lines representing the current distribution to overlaid grey shaded distributions from the previous 5 million year time slice), with an exception at 25 mya. During this time period, the primate body size distribution shifts rightward to larger sizes, corresponding to the increase in maximum size after the Grand Coupure species turnover in the fossil record.

**FIG. 4.**
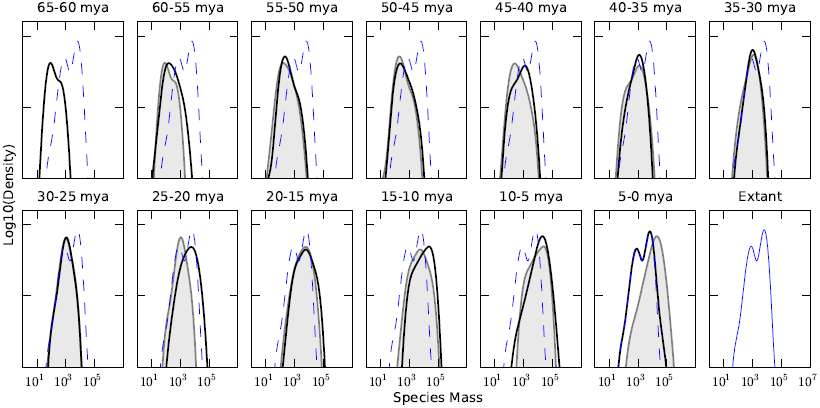
Primate body size distributions in 5 million year increments, from 65 mya to present. Each snapshot shows the distribution of species sizes in that period (solid black), the distribution in the preceding period (gray fill), and the extant distribution (blue online; dashed grey print). Over time there are two clear patterns: first, the primate body size distribution shifts rightward, towards larger sizes, and second, it becomes increasing left skewed.

When examining skewness in primate body size distributions using 5 million year snapshots, we see a gradual shift from a right-skewed distribution to the left-skewed distribution observed in extant primates (Figure 1). At 65 mya, there is a clear right skew to the body size distribution with a skewness of 0.20 ± 0.28 (Figure 5a). There are only gradual changes in the skewness of the distribution until 45 mya when the distribution skew becomes negative (−0.48 ± 0.37), and we observe a clear shift in the mean of the distribution to the right. This coincides with an increase in minimum body size. The left-skew trend in the distribution is robust through time from 45 mya to present (extant distribution skew −0.27 ± 0.11). The consistency in skewness after 45 mya suggests that the current left skew to the body size distribution is likely the result of internal gradual evolutionary changes and is not the result of recent anthropogenic extinction events or external forcing caused by specific environmental or climactic changes. Additionally, it would be unlikely under a null independent and identically distributed model with 0 skewness to observe skewness below 0 over multiple draws, as we do from 45 mya to present. This pattern in skewness over time suggests that the left skew of primate body size distributions is not accidental, and is instead the outcome of specific evolutionary pressures that lead to most species being relatively close to the largest size, while a few species are much smaller.

**FIG. 5.**
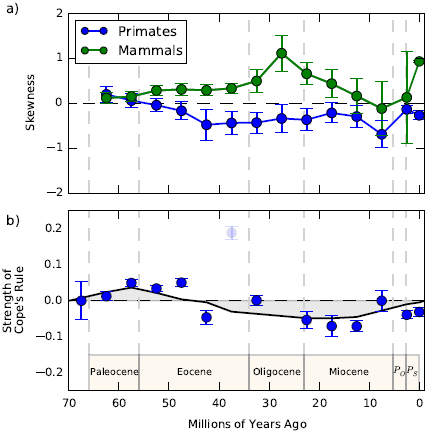
a) Body size distribution skewness, for primates (blue online; dark grey print) and North American terrestrial mammals (green online; light grey print) over 66 million years, in 5 million year increments. b) Strength of Cope’s rule for primates both in 5 million year increments (see text) and as a smoothed trend, representing the average of the four nearest estimates. (The estimate at 37:5 mya was omitted from the trend line calculation because it is based on only three ancestor-descendent pairs.)

The highly unusual evolution of primate body sizes is even more apparent when compared with patterns in body size distributions of terrestrial mammals in general (Figure 5a). The shift from positive to negative skewness for the primate distribution occurs 55 mya, falling well below the 0 skewness line 45 mya when the skewness becomes −0.48 ± 0.37 and remaining negative until present day. In contrast, the body size distribution of North American mammals is right skewed throughout time and highly right skewed from 55 to 15 mya. It is worthwhile to point out that the skewnesses of these two distributions are close 65 mya but diverge thereafter for 35 Ma. From 30 Ma onward, the distribution of North American mammals exhibits progressively less right-skewness, a trend whose beginning coincides with the Grande Coupure event, when global temperatures began to cool.

Variation in the skewness of body size distributions can be understood as the outcome of several macroevolutionary processes and constraints (Clauset et al. 2009), including Cope’s rule (Alroy 1998; Clauset and Redner 2009). For instance, cladogenetic variation in species sizes, in the context of increased extinction risk for large species and a physiological minimum size, has been shown to explain the right-skewed body size distribution (Clauset and Redner 2009) of terrestrial mammals. When descendant species are, on average, larger than their ancestor species (Cope’s rule), the right skewness created by the above processes will be enhanced (Alroy 1998). In contrast, if descendant species are, on average, smaller than their ancestor species (a negative value for the strength of Cope’s rule), the right skewness the above processes create will be reduced and possibly even converted into a left-skew distribution. Similarly, cladogenetic size variation in the presence of a maximum possible size and an extinction risk that increases as size decreases would naturally produce left-skewed size distributions, with Cope’s rule enhancing or mitigating that skewness depending on its direction, positive or negative. Examining the average strength of Cope’s rule over time for primates (Figure 5b), we observe an initial positive strength, which is concordant with most empirical investigations of Cope’s rule across clades and time. However, this measure of the average relative change in size from ancestor to descendent reverses sign at 47 mya, indicating that new primate species are typically smaller than their ancestors. This pattern remains consistent to present day.

This change in the direction of Cope’s rule could partially explain the left skewness observed in relatively recent and extant primate body size distributions, particularly if the extinction risk of primate species grew very weakly or not at all with increased size, However, if Cope’s rule alone fully accounted for the emergence of the left skew, we would expect the sign of the skewness to follow that of Cope’s rule after some delay or lag period. Instead, we observe the opposite pattern, in which the sign change in Cope’s rule follows the sign change in skewness, after a delay of 5 million years. This suggests that changes in the mechanisms governing the relative size of descendants to ancestors is not the primary cause of the unusual left-skew pattern among primates.

Instead, the observed pattern suggests the existence of a maximum body size for primates, around 230,000 g, which induces a left skew as the distribution presses up against the maximum. Just as a right-skewed distribution is the natural result of cladogenetic size variation in the presence of a hard minimum size and a slight correlation between extinction risk and body size (Clauset and Redner 2009; Shoemaker and Clauset 2014), a left skew would be a natural result of a hard upper bound on viable species size with an inverse or no relationship between size and extinction risk. In these circumstances, species are prevented from evolving past the maximum limit. The lower limit for terrestrial mammals is well understood to be due to thermoregulation constraints (Pearson 1948), which are unlikely to be the origin of a maximum size constraint.

We speculate that the maximum is related to an inability to meet the energetic and developmental needs of larger bodied but still highly encephalized, socially complex primates. Large brain size places greater energetic demands on individuals (Leonard et al. 2003), and the group social structure common among primates is well-established to be a function of brain size, specifically neocortical volume (Dunbar 1992). Maximum size would then be limited by the caloric intake necessary to maintain brain size and group structure, as diet quality decreases relative to body size in primates (Leonard and Robertson 1994). Indeed, a study of the diet of *Gigantopithecus*, an extinct primate genus (first appearance 5.33 mya and a final appearance in the fossil record 11, 700 years ago) containing the two largest primate species observed *(Gigantopithecus blacki* at 225, 000 g and *Gigantopithecus bilaspurensis* at 190, 000 g), suggests a dietary range insufficient to support its caloric needs through the climatic changes known to have affected its habitat (Bocherens et al. 2015). Thus, its large size and correspondingly large caloric requirements are hypothesized to have played a key role in its extinction (Bocherens et al. 2015). Furthermore, while all mammals exhibit decreased litter size and increased developmental time with increasing body size, these relationships are more severe for primates (Charnov and Berrigan 1993). Because primates have high encephalization coefficients, their offspring require longer developmental times compared to mammals more generally, limiting the number of offspring produced in a female’s lifetime. These constraints support our conjecture of a maximum size for primates, as larger species produce fewer offspring and have decreased diet quality, but still require enough energy to maintain high encephalization quotients and socially complex societies.

From the beginning of the Eocene to the end of the Oligocene, the size of the largest observed primate species is relatively stable, around 7000 g, well below the largest size of any extant primate (Figure 3a,b). This pattern suggests a transient maximum size for early primates, and it aligns well in time with the initial shift towards left skewness of the body size distribution and the beginning of New World primate radiation. Notably, soon after this initial maximum is reached, the clade’s size distribution begins exhibiting a negative skew. Such a lag is consistent with an initially right-skew distribution that encounters and then presses slowly up against a maximum limit. The subsequent release in the early Miocene of this transient limit is followed by a continued shift upward toward larger sizes, until, we speculate, a second maximum size limit is encountered, around 230, 000 g, in the mid-to-late Miocene. Throughout this period, the left-skew pattern continues, which suggests that the macroevolutionary forces that first created the left-skew pattern continued to shape the sizes of primates in this period.

During this time, the primate clade experienced a second radiation, this time in the New World. This event provides an unusual opportunity, in the form of a natural experiment, to investigate the relative contributions of external forcing events, such as climate change, from international evolutionary patterns, such as Cope’s rule in the evolution of body size and aggregate body size distributions. While Old World primates originated 66 mya, New World primates originated approximately 27 to 31 mya (Perez et al. 2013), providing two distinct radiation events within a single clade separated by time, location, and environmental conditions.

The rafting hypothesis is the leading explanation of the appearance of primates in the New World (De Oliveira et al. 2009). This method of travel would have a major impact on the initial size distribution of New World primates. In particular, rafting on small land or vegetative masses from the Old to New World would likely exclude many large primates from making the journey, biasing the initial distribution of New World primates towards the small end, which Figure 6 corroborates. From our dataset, the founder species for the New World radiation are indeed estimated to be between 600 and 3500 g, half as large as the maximum size observed in the Old World prior to 23 mya. Hind-casting using a radiation cone fitted to our database places the first appearance of New World primates at approximately 30 mya followed by a gradual increase in the maximum size of the distribution and a slower expansion towards smaller sizes.

**FIG. 6.**
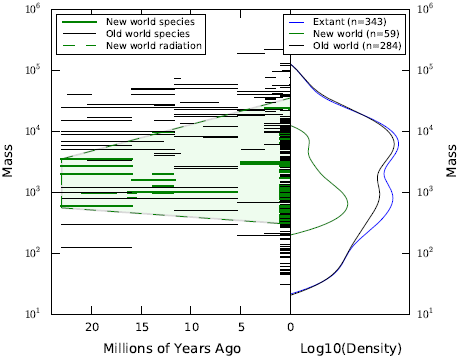
Left) Body sizes of all primate species from the first appearance of New World primate species to present, including a radiation cone (light green online; dashed grey print) for New World primates (green online; grey print) in comparison to all OldWorld species (black). Right) Smoothed extant body size distributions for all primates (blue online; dark grey print), New World primates (green online; light grey print), and Old World primates (black), illustrating the right-skewed pattern among NewWorld species and the left-skewed pattern among Old World species.

Independent phylogenetic and morphological analyses support the tracing of New World platyrrhine primate ancestry to one or several rafting events from the African continent in the late Eocene or early Oligocene (Kay et al. 2004). Stem platyrhinnes in South America, groups that branched off before the last common ancestor of modern platyrrhine primates, are geographically divided into three groups (Kay 2015). Of these, stem platyrhinnes found in the mid-latitudes represent the most ancient group, with *Branisella boliviana* dating back some 26 mya in what is now modern Bolivia (Takai and Anaya 1996). A second mainland stem clade of at least seven genera was found in higher latitudes, in areas of modern Chile and Argentina (between 34°and 52°S) between 21 mya and 16.5 mya (Kay 2015). The final stem clade, younger than the likely crown platyrrhine clade, is associated with the Greater Antilles on the islands of Cuba, Hispanola, and Jamaica from 18 mya and likely survived there until human occupation (MacPhee and Iturralde-Vinent 1995). Scant fossil evidence of species that may be part crown platyrrhines make identifying the exact origins of the clade difficult for both timing and location. However, fossil phylogenetic and molecular clock evidence both support origination at 20–24 mya (Hodgson et al. 2009; Kay 2015). Recently discovered fossil evidence from Panama indicates primates of the family Cebidae arrived by rafting from South America at 20.9 mya and therefore family-level diversification of extant platyrhinne families between 22 and 25 mya (Bloch et al. 2016).

After the initial founding event, the New World primate body size distribution expanded rapidly, with the maximum size increasing at a faster rate than the minimum size. The size disparity among New World primates shows no evidence of slowing, and we observe no evidence that the largest New World primates are close to the hypothesized maximum size observed in Old World primates. The extant distribution of body sizes for New World primates is also right skewed, similar to most other clades, including terrestrial mammals (Clauset and Erwin 2008), Equidae (Shoemaker and Clauset 2014), Cetacea Clauset (2013), and Old World primates 60 mya (Figure 6). The evolution of the Old and New World species body size distributions appear to have followed similar trajectories (comparing Old World from 66 mya to 45 mya and New World from 20 mya to present), with expansions of both maximum and minimum size as well as right-skewed distributions. One difference, however, is the length of expansion: the initial Old World primate radiation lasted approximately 10 million years while the New World primates expanded for 20 million years.

This contrast may be due to environmental differences, specifically the relative amount of grasslands versus tropical forests. The time period following the initial radiation of Old World primates is typified by high temperatures, tropical climates, and forests. Though considerable cooling took place during the Eocene, major losses of subtropical forests did not occur until near the Eocene-Oligocene boundary. Old World primates living during the Eocene therefore would have had habitats comprised primarily of forests with few open grasslands. Such ubiquity of trees suggests that a majority of primates would have led arboreal lives and would have been constrained in size by this lifestyle. In comparison, the time period surrounding the initial radiation of New World primates is marked by a slight increase in temperature and a small resurgence in subtropical forests followed by further temperature decreases and expanding grasslands. With more open space, advantages of larger body sizes would be pronounced. It is likely that this difference in environments resulted in a considerably smaller upper bound for Old World primates between 66 and 45 mya compared to New World primates.

The congruence of the pattern of expansion across these two primate radiations, despite the many environmental differences, suggests a common endogenous macroevolutionary process. Environmental variables appear to have only altered the length of primate radiations between the two and the transient maximum size for Old World primates. We speculate that were the New World radiation to continue, the size distribution would eventually become left skewed, like the Old World primates, as a result of pressing up against the social-complexity maximum size described above.

## IV. CONCLUSION

Investigating the evolutionary mechanisms that shape the distribution of species body sizes helps facilitate a deeper and more theoretically complete understanding of evolutionary dynamics. Body size distributions, particularly when studied over time, can also shed new light on the key factors that constrain and drive species sizes and on how related phenotypic traits coevolve with body size. Ultimately, understanding the evolutionary dynamics of species body sizes is a key part of developing more realistic evolutionary models.

After constructing a novel database of 742 primate species over the past 66 million years, we show that primates are not only unusual in their encephalization quotient, social structure, and ability to use tools, but also in the evolution of their body sizes. Unlike extant terrestrial mammals, birds, and lizards, as well as mammalian sub-clades like Equidae and Cetacea, which all exhibit a canonical right-skewed body size distribution, extant primate body sizes are left-skewed. This left skew is a robust pattern over the past 40 million years, and appeared only after a period of right-skewness that persisted throughout, and shortly after, the initial radiation. This highly unusual pattern can be understood as the natural outcome of a distribution first evolving up to and against a maximum size limit. This pattern is enhanced by a second unusual pattern: the tendency for descendant species to be smaller than their ancestors, which is the opposite of the more ubiquitous pattern of Cope’s rule, where descendants are typically larger than their ancestors.

These findings suggest that additional work is needed both to articulate evolutionary models of species body sizes (Clauset and Erwin 2008) that incorporate a maximum size parameter Gherardi et al. (2013), and models that can allow the directionality of Cope’s rule to change. The latter is particularly interesting, as the strength and direction of Cope’s rule is typically assumed to be fixed across evolutionary time and all members of a clade when modeling body size evolution.

Further, we showed how studies of body size distributions can elucidate taxonomic patterns, by comparing primate radiation with and without plesiadapiformes and diversity in body size after the gap in the fossil record during the Grand Coupure. By comparing New and Old World radiation events, we show that primate body size distributions initially mirror those of terrestrial mammals more generally and follow similar radiation patterns, suggesting internal consistency in the evolution of body size that can be modified by environmental factors. For primates, climate and the extent and prevalence of grasslands may have played a large role in shaping the speed of primate radiations and a maximum size through time. Investigating the precise mechanism by which these environmental factors shaped the distribution’s evolution would be valuable line of future work, and would shed new light on how broad-scale ecological processes shape the selective forces that drive clade-level body size dynamics.

Finally, our empirical findings support the belief that a relatively hard maximum species body size exists for primates, which we speculate is caused by the complex social structure of large primates, the large brains required for this lifestyle, and the lack of available energy from food to support larger brains. Circumstantial evidence from other studies suggests this mechanism is plausible, but further research is needed to determine its veracity. We look forward both to future investigations of primate body size evolution and its relationship to the many unusual characteristics of primates, and new body size evolution models that can better capture the full variability of body size distributions observed among animals.

## V. ACKNOWLEDGEMENTS

RCT was supported during this work by the University of Colorado’s Interdisciplinary Quantitative Biology program and NSF IGERT 1144807. LGS was supported by an NSF GRFP. We thank the University of Colorado’s Quantitative Think Tank for valuable discussions about the analyses.

